# Metal ion requirement for catalysis by 3′-5′ RNA polymerases

**DOI:** 10.1101/2025.03.21.644660

**Authors:** Brandon W.J. Iwaniec, Madison M. Allegretti, Jane E. Jackman

## Abstract

The two-metal ion mechanism for catalysis of RNA and DNA synthesis by 5′-3′ polymerases has been extensively characterized. The 3′-5′ polymerase family of enzymes, consisting of tRNA^His^ guanylyltransferase (Thg1) and Thg1-like proteins (TLPs), perform a similar nucleotide addition reaction, but in the reverse direction, adding Watson-Crick base paired NTPs to the 5’-ends of RNA substrates, yet the effect of divalent cations beyond magnesium has not been described. Here, we examined the effects of five divalent cations (Mg^2+^, Mn^2+^, Co^2+^, Ni^2+^ and Ca^2+^) on templated nucleotide addition activity and kinetics of 5’-activation by ATP catalyzed by recombinantly purified, metal-free TLPs from organisms from diverse domains of life. This work revealed that different TLPs exhibit distinct dependencies on the concentration and identity of divalent metal ions that support effective catalysis. The patterns of metal ion usage demonstrated here for TLPs evince features that are characteristic of both canonical 5’-3’ polymerases and DNA/RNA ligases. Similar to 5′-3′ polymerases, some metals were also seen to be mutagenic in the context of TLP catalysis. Furthermore, we provide the first direct evidence that both ATP and the NTP poised for nucleotidyl transfer are present in the active site during the 5’-adenylylation. These results provide the first in-depth study of the role of the two-metal ion mechanism in TLP catalysis that was first suggested by structures of these enzymes.

---

The tRNA^His^ guanylyltransferase (Thg1) family consists of the only known enzymes capable of catalyzing templated nucleic acid synthesis in the 3′-5′ direction(*1, 2*). This enzymatic function is performed in the opposite direction to canonical DNA and RNA polymerases. Thg1 enzymes, originally discovered in *Saccharomyces cerevisiae*, are restricted to eukaryotes and catalyze a non-templated 3’-5’ addition of a single nucleotide during the posttranscriptional incorporation of the essential 5’-guanosine (G_-1_) to tRNA^His^ (*3, 4*). The additional G_-1_ nucleotide distinguishes tRNA^His^ from all other tRNAs and allows for histidylation by histidyl-tRNA synthetase (*5–7*). The Thg1 family also contains homologs known as Thg1-Like Proteins (TLPs), which are found in all three domains of life and preferentially catalyze template-dependent 3′-5′ nucleotide incorporation (*8–10*). The biological functions of TLPs have proven more challenging to identify, with the *bona fide* physiological function of only two TLPs known to date, both in the slime mold *Dictyostelium discoideum*. These two enzymes each utilize template-dependent 3’-5’ RNA polymerase activity during maturation of several mitochondrial tRNA species (*9, 11*). For these reactions, TLPs catalyze template-dependent incorporation of nucleotides to the 5’-end of various tRNAs, using the nucleotides in the 3’-half of the tRNA as a template for Watson-Crick base pair selection (**Figure 1A**). While the biological functions of most TLPs remain to be defined, extensive *in vitro* biochemical characterization has revealed that 3’-5’ RNA polymerase activity is a shared biochemical feature of TLP enzymes from all domains of life (*1, 8*).

**FIGURE 1:**
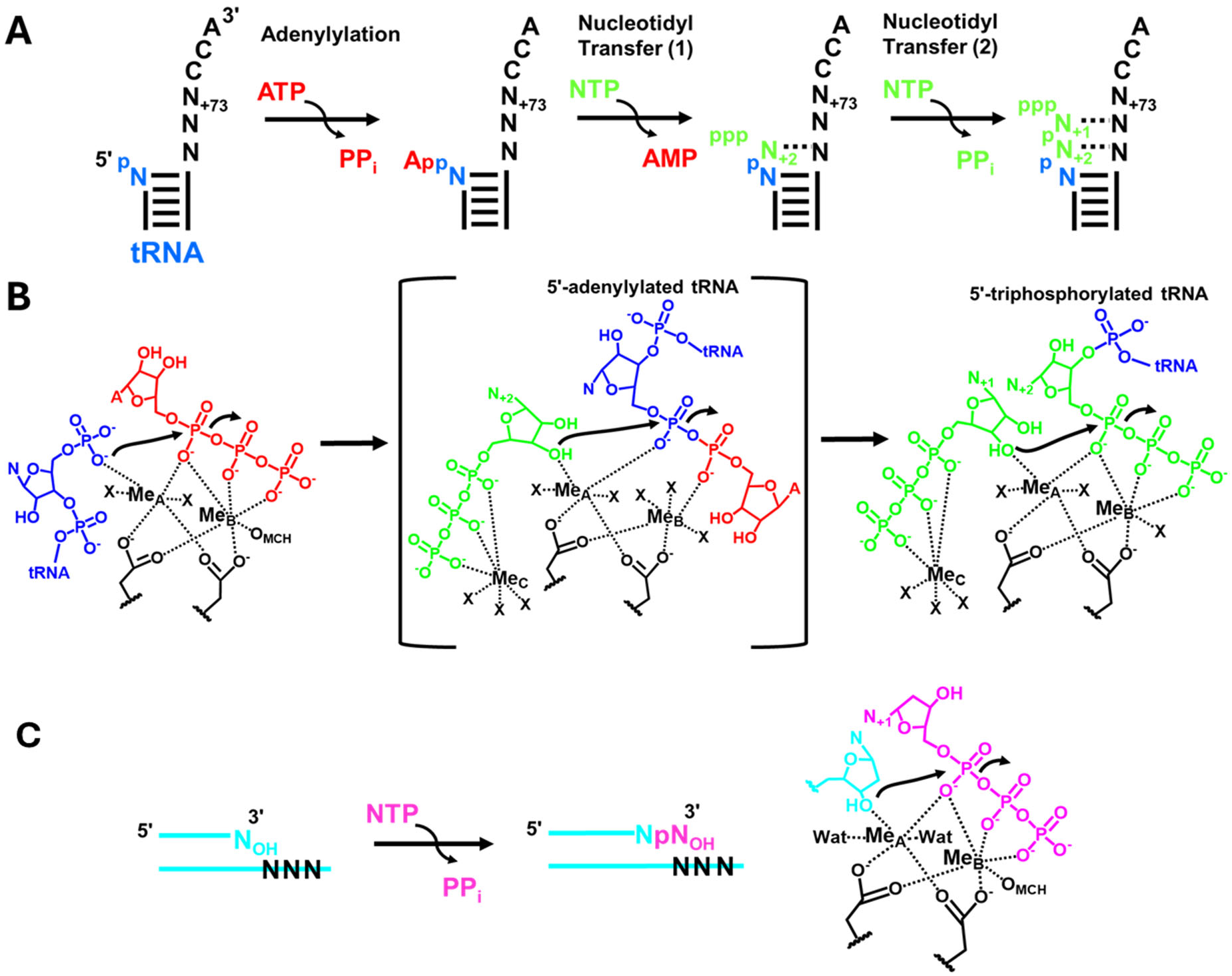
The two metal ion mechanism of 3’-5’ and 5’-3’ polymerases. **(A)** Schematic of 3’-5’ polymerase activity during tRNA 5’-end repair. A 5’-truncated monophosphorylated tRNA (blue) is undergoes three distinct catalytic steps to restore multiple missing 5’-nucleotides. Each added NTP (green) is selected to create a Watson-Crick base pair, using the nucleotides on the 3’-end of the tRNA (black) as a template. For the substrate depicted here, TLPs add the N_+2_ nucleotide to a 5’-adenylylated tRNA intermediate, then the N_+1_ nucleotide (green) is added to the 5’-triphosphorylated tRNA that is generated after the first N_+2_ addition. **(B)** Known and proposed roles of metal ions in the distinct steps of 3’-5’ polymerase activity during tRNA 5’-end repair. The two metal ion mechanisms for adenylylation and nucleotidyl transfer (2) shown are based on available structures, but no structure is available to show the enzyme poised to catalyze nucleotidyl transfer (1) (as indicated by brackets). The cartoons are meant to highlight groups that are known or proposed to coordinate each metal and do not accurately replicate actual coordination geometry; X residues indicate waters that are observed or proposed based on available structures. **(C)** Schematic of 5’-3’ polymerase activity of a representative DNA polymerase adding an NTP (magenta) to the 3’-end of an elongating primer (teal). The two metal ion mechanism involving attack of the 3’-hydroxyl on a 5’-triphosphate is directly analogous to nucleotidyl transfer (2), with the only difference being the orientation of the attacking nucleophile from elongating DNA (for 5’-3’ polymerases) vs. from the incoming NTP (for 3’-5’ polymerases).

Due to their novel enzyme chemistry, it was not necessarily surprising that the first structures of Thg1 and TLP enzymes revealed a unique overall protein fold (*12, 13*). However, these structures also revealed a striking local similarity between the active site of Thg1/TLP enzymes and the catalytic palm domain of canonical 5’-3’ DNA polymerases such as T7 DNA polymerase and bacterial DNA polymerase II(*12*). A superposition of the catalytic sites of human Thg1 and T7 DNA polymerase revealed similar positioning of conserved carboxylate-containing residues that coordinate two divalent metal ions in both proteins, suggesting an overall similarity in mechanism between Thg1/TLP enzymes and the well-studied canonical DNA and RNA polymerases (*12*) (**Figure 1B**). Alteration of metal-coordinating amino acids in several different Thg1/TLP 3’-5’ polymerases to alanine severely reduces or eliminates enzyme activity, consistent with similar use of two-metal ion catalysis by Thg1 and TLPs (*12, 14*). The ability of Thg1/TLPs to use the two-metal ion site to catalyze the same overall chemistry but in a different direction (3′-5′ vs. 5′-3′) is linked to the different orientation of the incoming nucleotide and nucleic acid substrates in these enzymes, but the molecular features that enable the transitions that occur around the metal ions and enable its distinct chemistry still remain incompletely understood.

The two metal ion mechanism has been studied extensively in the context of canonical 5’-3’ synthesis of DNA and RNA. In 5’-3’ polymerases, one metal ion coordinates the α-phosphate of the incoming dNTP and the 3′-OH of the primer strand, thus decreasing the pKa of the elongating primer hydroxyl (**Figure 1C**). The second metal ion coordinates phosphate oxygens of the incoming dNTP, which helps to neutralize the negative charge of the developing transition state while assisting with the departure of pyrophosphate after the new nucleotide has been added. Thg1/TLP enzymes catalyze a directly analogous nucleotidyl transfer reaction during 3’-5’ polymerization, with an attacking 3’-OH (from the incoming NTP) acting on the α-phosphate of a 5’-triphosphorylated RNA (**Figure 1**, Nucleotidyl Transfer (2)). However, Thg1/TLP enzymes are also capable of initiating 3’-5’ polymerization from a 5’-monophosporylated RNA end using a 5’-activation step (**Figure 1**, Adenylylation). During this reaction, the two metal ion site is used to activate the 5′-monophosphorylated end of the RNA and create a 5′-adenylylated (or in some cases guanylylated) intermediate (**Figure 1B**). Subsequently, the first nucleotidyl transfer step occurs where the 3′-OH of the incoming NTP attacks the adenylylated intermediate (**Figure 1**, Nucleotidyl Transfer (1)). After one NTP has been added to the growing RNA to initiate repair, the product retains the 5’-triphosphate from the first added nucleotide, and therefore the next and all subsequent NTPs are incorporated by an ATP-independent mechanism that only requires nucleotidyl transfer chemistry, directly analogous to the polymerase chemistry catalyzed by 5’-3’ polymerases (**Figure 1**).

The RNA activation step (adenylylation/guanylylation) is a distinct feature of 3’-5’ polymerases, since it is only required during the first round of NTP addition to a monophosphorylated RNA (*4, 14, 15*). The step is utilized for at least one biological substrate, since Ribonuclease P generates the 5’-monophosphorylated tRNA^His^ that is the substrate for Thg1 during tRNA^His^ maturation in eukaryotes (*3, 16, 17*). Most of the existing structures of eukaryotic, bacterial and archaeal 3’-5’ polymerases show the active site poised to carry out this adenylylation step, with a bound ATP or GTP in the activation site (*12, 13, 18, 19*). One structure of an archaeal TLP active site with 5’-triphosphorylated tRNA and non-hydrolyzable NTP suggests the involvement of a third metal in positioning the incoming NTP that is consistent with prior kinetic characterization (*20*). However, none of these available structures provide molecular details about the multiple transitions of reactive groups that must occur around the metal ions as the reaction progresses between these steps. Moreover, the specific nature of the metal ion requirements for catalysis by 3’-5’ polymerases have not been characterized.

In this work, we characterized the metal ion dependence of several members of the 3’-5’ RNA polymerase family representing all three domains of life. Since TLPs generally exhibit more flexible RNA substrate recognition than their Thg1 counterparts and therefore are excellent candidates for engineering to exploit this unusual activity, this investigation was focused on TLP family members (*1, 21, 22*). Catalytic activities of purified metal-free enzymes were compared in the presence of different concentrations of divalent cations representing a range of sizes and electronic properties. We determined that 3’-5’ polymerases generally demonstrate similar metal ion requirements for overall catalysis to those observed with most 5’-3’ polymerases, with high activity observed in the presence of Mg^2+^, Mn^2+^, and Co^2+^, and to a significantly lesser extent, Ni^2+^. Surprisingly, however, Ca^2+^ is able to support the chemistry of the 5’-adenylylation reaction that occurs with some TLP substrates. This is an intriguing feature given that the overall arrangement of the two-metal ion active site appears to be similar for both reactions and reveals a key mechanistic difference between 5’-3’ vs. 3’-5’ polymerases. The ability of Ca^2+^ to uniquely catalyze the activation step enabled us to also provide the first direct experimental evidence that the fidelity of base-pair selection is promoted by the presence of two NTPs (one for 5’-end activation and one for nucleotidyl transfer) that simultaneously occupy the active site during the first (adenylation) step of the reaction.

## Materials and Methods

### Nucleotide and Reagents

NTPs (100 mM LiCl salts) used for enzyme assays were purchased from Roche; [γ-^32^P] ATP (6000 Ci/mmol) was purchased from Revvity. Oligonucleotides were purchased from Sigma. Enzymes used for labeled tRNA substrate preparation (T4 polynucleotide kinase (PNK) and Quick calf intestinal phosphatase (CIP)) were from New England Biolabs. For TLP kinetic assays, ribonuclease A was purchased from Ambion, and calf intestinal alkaline phosphatase (CIAP) was purchased from Invitrogen.

### Recombinant Protein Expression and Purification

Plasmids for T7 RNA polymerase-dependent expression of MxTLP (UniProt KB Q1CZS0), AcaTLP2 (UniProt KB J9XZJ1, MsTLP (UniParc UPI000B984832), BtTLP (UniProt KB A0AAN4HKC4), and DdiTLP4 (UniProt KB Q86IE7) were transformed into *E. coli* strain BL-21 DE3 (pLysS). Cultures were grown at 37°C in 1L of LB media containing 100 µg/µL ampicillin until OD_600_ reached 0.4-0.6, induced by addition of final concentration of 1 mM isopropyl β-D-galactopyranoside. Cultures were induced at 18°C for ∼20 hours (MxTLP, AcaTLP2, and MsTLP) or induced at 37°C for 5 hours (BtTLP and DdiTLP4), harvested at 5000 x g for 20 min at 4°C, resuspended in 20 mL of H_2_O, divided in half, and centrifuged at 5000 x g for 20 min at 4°C to pellet cells. Each cell pellet corresponding to 0.5 L culture was resuspended in 20 mL (for sonication) or 10 mL (for bead beater homogenization) of resuspension buffer (20 mM HEPES, pH 7.5, 1 M NaCl, 0.5 mM β-mercaptoethanol) containing 1 mg/mL lysozyme and protease inhibitors (1 µg/µL pepstatin, 1 µg/µL leupeptin, and 1 mM phenylmethylsulfonyl fluoride (PMSF)). Crude extract was obtained either by sonication (for MxTLP) or bead beater (for all other proteins). Sonication of the mixture was performed with alternating cycles of 4 sec on/6 sec off at 55% amplitude for 5 min total sonication, incubated on ice for 5 min, and repeated 2 more times. Bead beater homogenization of the mixture was done using alternating cycles of 1 min homogenization/2 min on ice using a Mini-BeadBeater 16 (Biospec Products), for a total of 3 cycles. The crude extract from either lysis method was centrifuged at 5000x g for 25 min to remove debris. Extracts were diluted with an equal volume of a no salt buffer lacking NaCl (20 mM HEPES, pH 7.5, 0.5 mM β-mercaptoethanol; resulting in 0.5M NaCl total) and mixed for 30 minutes with 1 mL of Talon resin that had been pre-washed in 0.5 M NaCl buffer (20 mM HEPES, pH 7.5, 0.5 M NaCl, 0.5 mM β-mercaptoethanol). The resin was washed with 20 volumes of 0.5 M NaCl buffer two times, followed by 15 volumes of 0.5 M NaCl with added 10 mM imidazole three times. Each wash step consisted of 10 min mixing followed by low-speed centrifugation to pellet resin prior to removal of wash buffer. The resin was packed into a column and rinsed with one column volume of 0.5M NaCl/10 mM imidazole buffer, and proteins were eluted using 0.5 M NaCl buffer plus 250 mM imidazole. Peak protein fractions as determined by Bio-Rad protein assay were pooled.

### Metal-Free TLP purification

Apo-TLPs were purified by adding EDTA (final concentration 100 mM) to the eluted enzyme-containing pooled fractions, then incubating on ice for 30 minutes, mixing gently every 10 minutes. Removal of EDTA and concentration of the TLPs was done using successive cycles of dilution into EDTA and metal-free buffer (20 mM Tris, pH 7.5, and 0.5 M NaCl) followed by concentration using Amicon® Ultra-4 Centrifugal Filters. Once the concentration of EDTA was below 50 nM (based on serial dilution calculations), the TLPs were flash frozen on dry ice and stored at −80°C. Proteins were judged to be ≥90% pure based on visual inspection of the of the purified enzyme preparation using sodium dodecyl sulfate polyacrylamide gel electrophoresis (PAGE). Concentrations of the proteins were determined using Bio-Rad protein.

### *In Vitro* Transcription and 5′-end labeling of tRNA

*In vitro* transcription reactions were performed with plasmids that contained a T7 promoter sequence upstream of the indicated tRNA gene and a BstNI site for linearization of the plasmid downstream of the tRNA gene that creates the tRNA 3’-CCA end during runoff transcription. Transcription was performed at 37°C for 2 hours in reaction mixtures containing 50 mM Tris-HCl (pH 8.0), 1 mM spermidine, 30 mM MgCl_2_, 8 mM DTT, 2 mM of each NTP, 0.1-0.2 mg/mL template DNA, and 10-50 µg T7 RNAP. Reactions were incubated at 37°C for 30 min with 0.06 U/µL of DNase I. Triphosphate tRNAs create via *in vitro* transcription were PAGE purified using 10% polyacrylamide, 4 M urea gels, followed by phenol/chloroform/isoamyl alcohol extraction and ethanol precipitation. To create 5′-[^32^P] monophosphorylated tRNA, 40 pmol of each purified transcript were treated with 20 units of Quick CIP at 37°C for 15 min to remove all 5’-phosphates, followed by treatment with 2 units of T4 PNK and 100 µCi [γ-^32^P] ATP at 37°C for 30 min. 5′-[^32^P]-tRNAs were PAGE purified using 10% polyacrylamide, 4 M urea gels, followed by phenol/chloroform/isoamyl alcohol extraction and ethanol precipitation.

### Phosphatase Protection Assay for 3′-5′ tRNA Repair

The 5′-[^32^P]-tRNAs were used as the substrate in a 3’-5’ RNA repair assay where addition of the missing G_+2_ nucleotide could be visualized using a phosphatase protection assay described previously (*14*). Briefly, reactions contained 25 mM HEPES, pH 8, 125 mM NaCl, 0.1 mM ATP, 1 mM GTP, and reactions were started by initiation of 10 µM purified TLP enzyme. Divalent cation titration assays contained 0.25, 0.5, 1, or 2 mM of the indicated metal chloride (MgCl_2_, MnCl_2_, CoCl_2_, CaCl_2_, or NiCl_2_). After incubation for the indicated time points (1-30 min) at room temperature, reactions were quenched by removal of an aliquot (5 µL) to a tube containing 0.5 µL of 10 µg/µL RNase A and 0.5 µL of 0.5 M EDTA, pH 8.0, followed by digestion for at 50°C for 10 min, and treatment with 0.5 U CIAP in 1x reaction buffer at 37°C for 30 min. Reactions were spotted (2 µL) on silica-TLC plates and resolved in an n-propanol:NH_4_OH:H_2_O (55:35:10 v:v:v) solvent system. The TLC plates were visualized and quantified using PhosphorImager and ImageQuantTL software.

### Adenylylation Assays

Single turnover enzyme assays were performed to determine kinetic parameters for the adenylation step of 3′-5′ tRNA repair using the same phosphatase protection assay described above, except to isolate the adenylylation step, GTP was not added to the reaction (*14*). The k_obs_ for each condition was determined by plotting time courses of product formation, which were plotted and fit to a single exponential rate equation (eq. 1) using Kaleidagraph (Synergy software), where Pt is the fraction product formed at each time and ΔP is the maximal amount of product conversion observed during each time course. All reported rates are determined from three independent replicates, unless otherwise indicated.

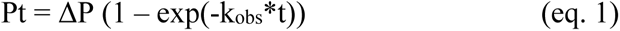

For reactions that occurred with rates that were too slow to complete the reaction progress curve during measured time points, the method of linear initial rates was used to estimate k_obs_, using the linear initial slope of the progress curve (v_o_) and ΔP determined in a separate assay using the same substrate, and fitting to eq. 2.

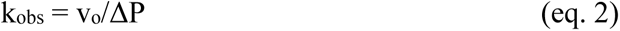

## Results

### Investigation of the divalent cation requirement for TLP 3’-5’ polymerase activity

Based on the assumed two-metal ion mechanism first proposed from Thg1/TLP structures, biochemical characterization of TLPs to date has been performed consistently in the presence of 10 mM Mg^2+^. To determine whether other divalent cations could support 3’-5’ polymerase chemistry, we first sought to produce metal-free apoenzyme using an alternative purification strategy for recombinantly expressed TLP enzymes in *E. coli*. Briefly, we omitted MgCl_2_ from all protein purification buffers used during affinity purification and then incubated the eluted pooled protein fractions in EDTA (100 mM final concentration) to remove any remaining metals from the enzyme preparation. Buffer exchange was performed by successive dilution/concentration in EDTA-free buffer until [EDTA] < 50 nM, corresponding to∼1/10,000 – 1/20,000 of the final [TLP]. To test activity of the purified metal-free proteins, we utilized an *in vitro* phosphatase protection assay for 3’-5’ addition that has been used extensively to characterize 5’-end repair reactions like the one shown in Figure 1A. The substrate used here was a 5’-truncated *D. discoideum* mitochondrial tRNA^Ile^ transcript that is missing only one nucleotide (G_+1_) to be added opposite nucleotide C_72_ to complete the aminoacyl-acceptor stem of the tRNA, thus simplifying the number of reaction products to be detected (*11, 15*) (**Figure 2A**). The products of the activation (AppC) and nucleotidyl transfer (GpC) steps have been extensively characterized and validated with this tRNA(*10, 15*). Although this mt-tRNA^Ile^ is one of the biologically relevant substrates for the *D. discoideum* enzyme (DdiTLP3) that repairs tRNA *in vivo*, it is also an efficient substrate for *in vitro* 5’-end repair by other TLPs, even though many of these enzymes are not believed to utilize similar substrates *in vivo* in their relevant organisms.

**FIGURE 2.**
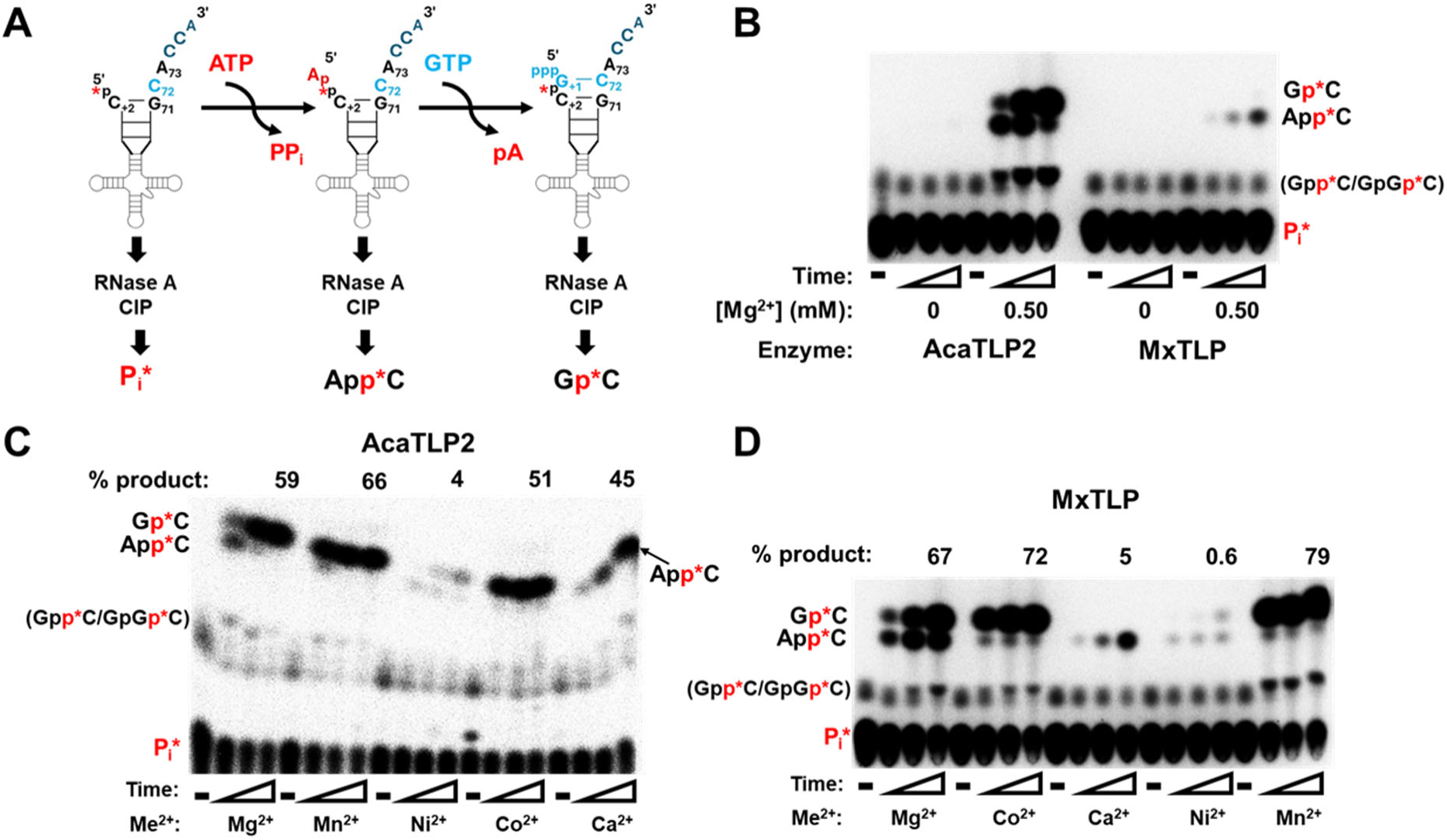
Divalent metal ions support 3’-5’ polymerase activity of diverse TLPs. **(A)** Schematic of the phosphatase protection assay used to measure 3′-5′ polymerase activity to add the single missing nucleotide (G_+1_) to the 5′-end of 5′-^32^P-mitochondrial-tRNA^Ile^ from *D. discoideum*. The identity of the radiolabeled products formed from the unreacted substrate (P*_i_), 5’-adenylylated intermediate (App*C) and 5’-end repaired product (Gp*C) after treatment with RNase A and CIP are shown below each species. These products are resolved by thin-layer chromatography (TLC), as indicated on panels B-D. **(B)** Phosphatase protection assay results with AcaTLP2 and MxTLP purified in the absence of Mg^2+^ (as described in Materials and Methods). Assays contained 10 µM enzyme, 1 mM GTP, 0.1 mM ATP, 5′-^32^P-mitochondrial-tRNA^Ile^, and Mg^2+^ (as indicated). No enzyme control lanes are indicated with “-”. Triangles indicate increasing amount of reaction time, with 1, 5 and 30 min time points shown for each reaction condition. In the reactions with AcaTLP2, an additional lower-migrating product is also observed, which could correspond to two possible products that have been observed previously with other TLPs and are the result of 5’-activation with GTP instead of ATP (Gpp*C) or addition of a second non-templated GTP nucleotide (using the nucleotidyl transfer (2) mechanism shown in Figure 1A and generating GpGp*C). These two products do not resolve well on this TLC system and therefore cannot be conclusively identified which is shown here. **(C and D)** Representative phosphatase protection assays comparing enzymatic activity of apo-AcaTLP2 **(C)** and apo-MxTLP **(D)** with various divalent cations as indicated (2 mM each). Assays contained 10 µM of either enzyme, 1 mM GTP, 0.1 mM ATP, 5′-^32^P-mitochondrial-tRNA^Ile^. No-enzyme controls lanes are indicated with “-”.Reaction products observed at 1, 5 and 30 min were resolved by TLC. Total % reaction products formed at the 30 minute time points with each metal ion were quantified and are indicated above each set of reactions. Note that the anomalous mobility of the products for the reactions shown in panel **(C)** causes a distinct pattern for the App*C product generated in the presence of Ca^2+^, but the identity of this product has been validated in other assays.

We used the metal-free protocol to purify two TLP enzymes from different domains of life (a eukaryotic amoebozoan *Acanthamoeba castellanii* (AcaTLP2) and bacterial *Myxococcus xanthus* (MxTLP)) and demonstrated that both enzymes lacked any detectable catalytic activity in the absence of added metal (**Figure 2B**). Upon addition of 0.5 mM Mg^2+^, AcaTLP2 exhibited robust G_+1_-addition activity (corresponding to Gp*C), similar to enzyme purified and tested under standard conditions in earlier studies, with a smaller amount of 5’-adenylylated intermediate (corresponding to App*C) remaining in the assay at the longest (30 min) time point (**Figure 2B**). However, MxTLP exhibited relatively little product formation in the presence of 0.5 mM Mg^2+^, with only a small amount of the 5’-adenylylated tRNA intermediate generated at the 30 min timepoint and little, if any, detectable G_+1_ addition. To determine whether higher concentrations of metal could increase the activity of MxTLP, each apoenzyme was tested in the presence of increasing [Mg^2+^], ranging from 0.25 to 2 mM. MxTLP activity was strongly dependent on the concentration of added Mg^2+^ across this range, and at 2 mM Mg^2+^ MxTLP exhibited similar robust formation of G_+1_-addition product to AcaTLP2 (**Figure 3A**). AcaTLP2 activity only showed a modest increase in activity across the same Mg^2+^ concentration range (**Figure 3B**).

**FIGURE 3.**
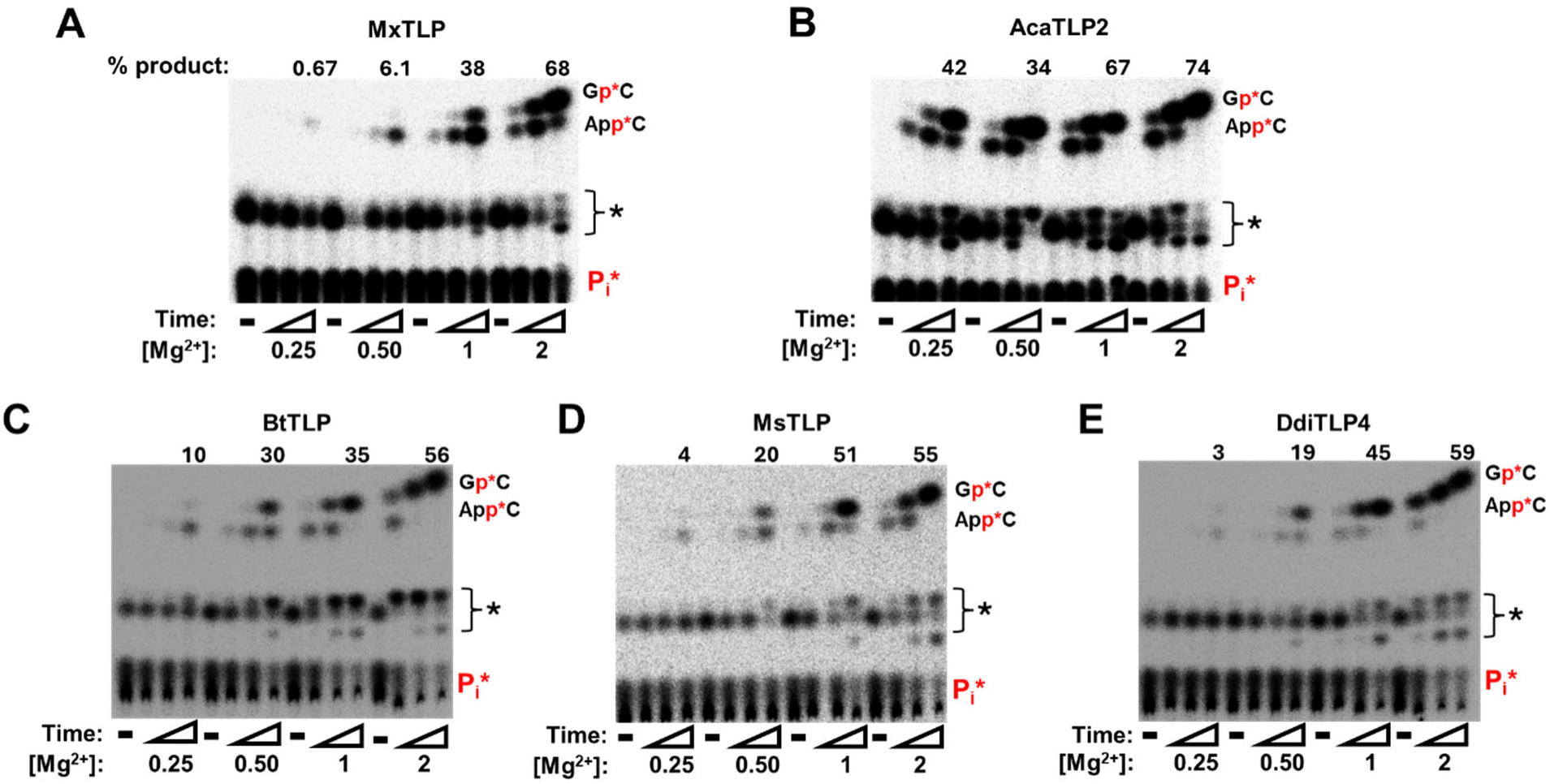
AcaTLP2 exhibits a distinct pattern of activity with [Mg^2+^] compared to four other TLPs. Phosphatase protection assays were performed with 5′-^32^P-mitochondrial-tRNA^Ile^ with 10 µM MxTLP **(A)**, AcaTLP2 **(B)**, BtTLP **(C)**, MsTLP **(D)**, or DdiTLP4 **(E)**, 1 mM GTP, 0.1 mM ATP, and 0.25 – 2 mM [Mg^2+^], as indicated below each figure. No enzyme controls lanes are indicated with “-”. Timepoints corresponding to 1, 5, and 30 min reactions are shown for each reaction, and the indicated products were resolved by TLC. The total % product indicated above each set of reactions is for the 30 min time point at each [Mg^2+^].

At higher concentrations of Mg^2+^, we also noted time-dependent production of lower migrating products, which have also been observed previously in phosphatase protection assays(*10, 15*). These products are known to arise from at least two additional reactions that are possible with these substrates, one of which is the alternative use of GTP as a 5’-end activating nucleotide, indicated by Gpp*C in these assays, and the other corresponds to the addition of a second non-templated GTP addition after incorporation of G_+1_, indicated by GpGp*C. These two products are not typically well-resolved in this TLC system, and often co-migrate, making it difficult to conclusively determine whether one or both of these products are made in most assays. Nonetheless, since for most of these enzymes (with the possible exception of BtTLP), these are minority reaction products, we chose to focus on quantification of the well-resolved and validated 5’-adenylylate and G_+1_ addition products to compare reactivities in these and subsequent analyses.

We then tested 3’-5’ nucleotide addition activity of AcaTLP2 and MxTLP in the presence of 2 mM of each of five divalent metals (Mg^2+^, Mn^2+^, Co^2+^, Ni^2+^, and Ca^2+^) (**Figure 2C and D**). With Mg^2+^, Mn^2+^ and Co^2+^, both TLPs created similarly high amounts of total product (51-79%) at the longest (30 min) time point. Interestingly however, accumulation of the 5’-adenylylated intermediate was notably lower in reactions with Co^2+^ and Mn^2+^ than in the Mg^2+^-containing reactions, especially for MxTLP (**Figure 2D**), with a larger fraction of the total reaction products in Co^2+^ and Mn^2+^ corresponding to G_+1_ addition at the longest time points. These data suggest that the ability of each enzyme to progress through step 2 (nucleotidyl transfer) with Mg^2+^ is not as robust as it is in Mn^2+^ and Co^2+^. Reactions performed in the presence of Ni^2+^ were also able to progress through both steps to create G_+1_ addition product, but at a severely reduced overall catalytic efficiency (creating 4% and 0.6% total product in AcaTLP2 and MxTLP, respectively). Finally, no G_+1_ addition product was observed in Ca^2+^-containing reactions for either enzyme, although 5’-adenylylated intermediate was readily detected in the presence of Ca^2+^.

### Distinct metal concentration dependencies exhibited by TLPs

Since we observed MxTLP and AcaTLP2 to vary in their dependence on metal ion concentration (**Figure 3A and B**), we tested the generality of these trends with other TLPs. tRNA repair activity was measured as a function of [MgCl_2_] for three other TLPs representing enzymes from all three domains of life (*Bacillus thuringiensis* TLP (BtTLP), *D. discoideum* TLP4 (DdiTLP4), and *Methanobrevibacter smithii* TLP (MsTLP)). A comparison of product formation revealed that BtTLP, DdiTLP4 and MsTLP all exhibited a similar pattern to MxTLP; catalytic activity increases substantially across the tested Mg^2+^ concentration range (0.25 – 2 mM) (**Figure 3C-E** and **Table 1**). Thus, the ability of AcaTLP2 to efficiently catalyze nucleotide addition even at low (0.25 mM) [Mg^2+^] is likely a unique mechanistic feature of this homolog, which is consistent with other unusual enzyme characteristics recently demonstrated for AcaTLP2 (*22*).

**Table 1:**
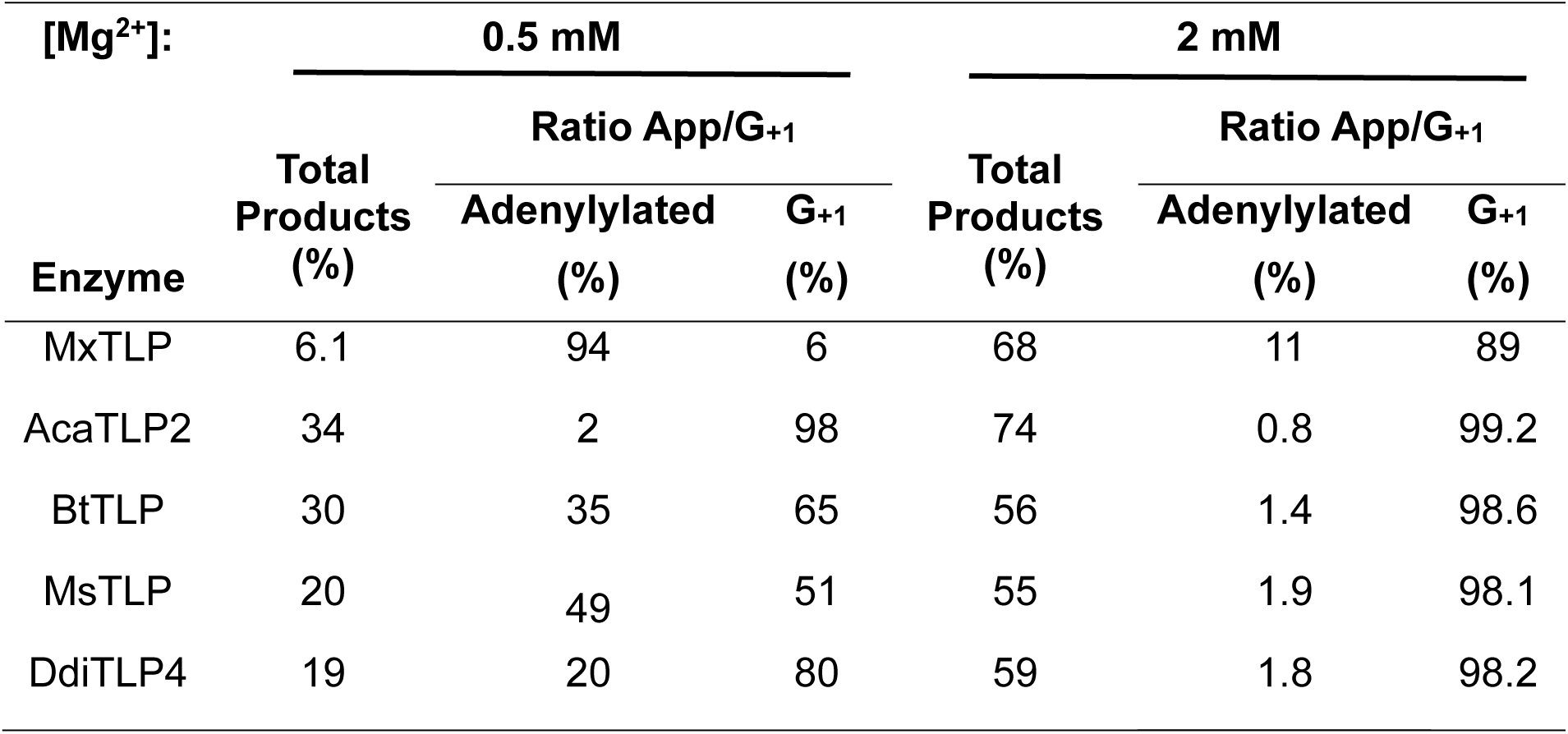
Quantification of 3’-5’ addition reactions with diverse TLPs in Mg^2+^.

Quantification of these reactions revealed distinct ratios of adenylated (step 1) vs. nucleotidyl transfer (step 2) products between high vs. low [Mg^2+^] that suggests the impact of metals on the two steps of catalysis differs among the tested enzymes (**Table 1**). At high [Mg^2+^] (2 mM), all five enzymes produced relatively high amounts of total product after 30 min, ranging from 55% with MsTLP to 74% with AcaTLP2. The fully repaired (G_+1_ addition) products predominate with all five enzymes at high (2 mM) Mg^2+^, reaching between 89-99% of the total reaction products at the 30 min time point. However, at lower concentrations of Mg^2+^, significant differences are observed. For example, at 0.5 mM Mg^2+^ where activity is readily detectable for all five tested enzymes, 98% of the total products produced by AcaTLP2 correspond to the fully repaired G_+1_ tRNA, while for MxTLP this ratio is almost reversed, with 94% of the total reaction products corresponding to the 5’-adenylylated intermediate. The other tested enzymes show more intermediate effects at low Mg^2+^, with 50-80% of total products corresponding to G_+1_ addition in these assays. Thus, AcaTLP2 is able to efficiently progress through both steps of the G_+1_ addition reaction even at low concentration of metal, while other enzymes accumulate more 5’-adenylated intermediate at lower [Mg^2+^].

### Kinetics of 5’-end activation reveal distinct Mg^2+^-dependent trends

Since 5’-activation and nucleotidyl transfer seem to depend differently on metal concentration, we sought to further quantify these effects. We utilized a single turnover kinetic assay that we previously developed to measure the rate of 5’-adenylylation (*14*). Here, k_obs_ was measured using the phosphatase protection assay under single turnover conditions ([E] >> [tRNA]) in the presence of saturating concentrations of ATP (0.1 mM) and Mg^2+^(2 mM) (**Figure 4A**). The omission of the correct Watson-Crick base pairing GTP substrate for this 5’-[^32^P]-tRNA^Ile^ substrate is intended to stop the reaction before nucleotidyl transfer could proceed through step 2 to create G_+1_ product with this tRNA (compare with reaction schematic shown in **Figure 2A**). Nonetheless, AcaTLP2 proceeded through nucleotidyl transfer to add a non-Watson-Crick A_+1_ to the adenylated tRNA (**Figure 4B**). Non-templated ATP addition in the absence of the correct Watson-Crick base pairing NTP (GTP in this case) has been observed for both Thg1 and TLP-type enzymes (*2, 10, 15, 23*). Interestingly, MxTLP did not form any of the A_+1_ nucleotidyl transfer product at any concentration of Mg^2+^ despite reaching similar amounts of 5’-adenylylated intermediate. Since 5’-adenylylation is rate-determining under these single turnover conditions, both products (AppC and A_+1_pC) were used to determine k_obs_ by fitting to the single-exponential rate equation (eq. 1) (**Figure 4C**, **Table 2**).

**FIGURE 4.**
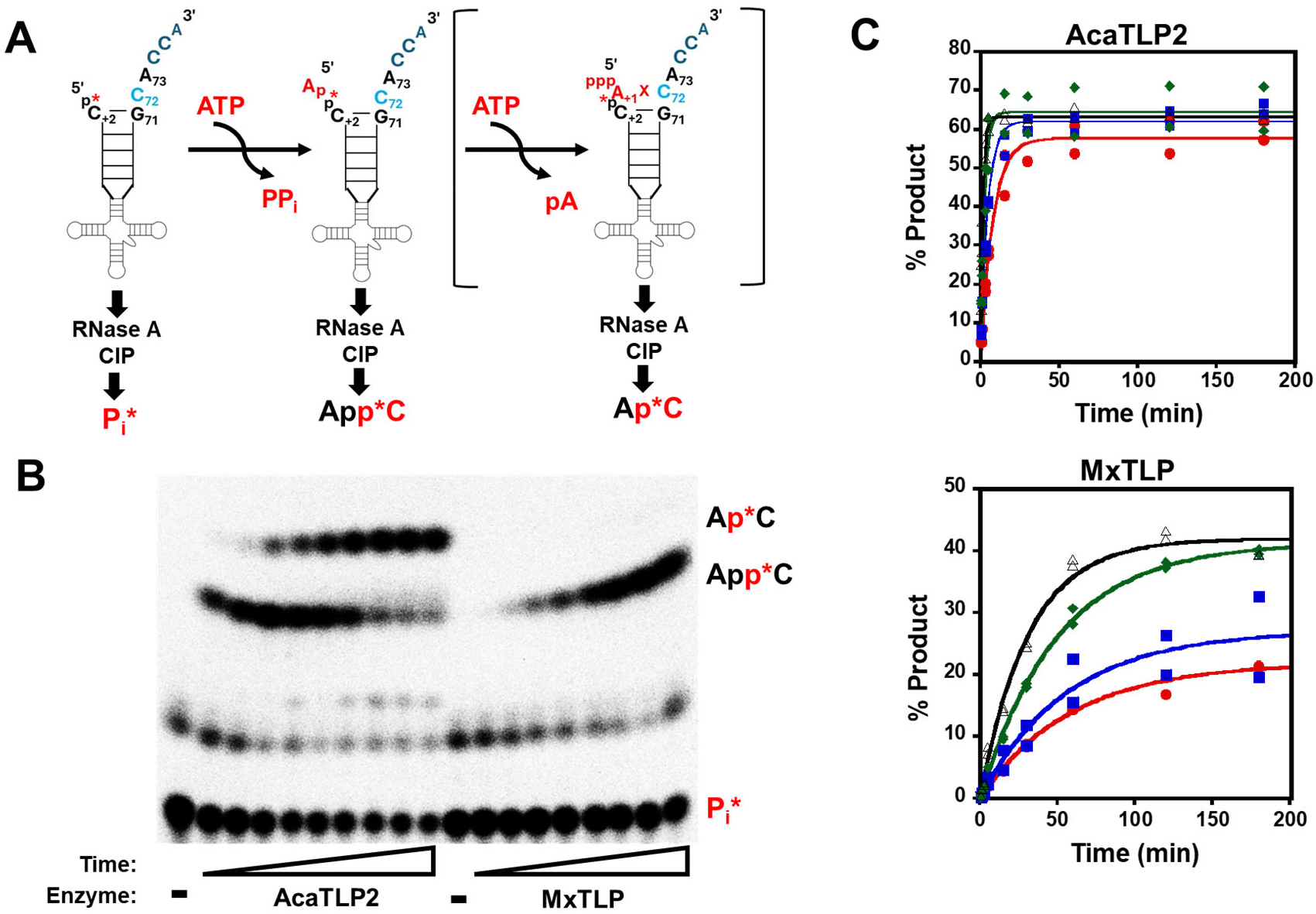
Effects of Mg^2+^ concentration on the rate of 5’-adenylylation catalyzed by TLPs. **(A)** Schematic of the 5’-adenylylation assays used with the 5′-^32^P-mitochondrial-tRNA^Ile^ substrate to measure single turnover rates. The phosphatase protection assay is performed as described for the complete reaction, except that only ATP is included in the reactions. Under these conditions, enzymes accumulate 5’-adenylate (App*C), but some TLPs are capable of progressing through a non-templated nucleotidyl transfer step with varied efficiency (shown in square brackets) to generate the 5’-A_+1_ reaction product (Ap*C). **(B)** Representative phosphate protection assays containing 10 µM of the indicated enzyme, 0.1 mM ATP, 2 mM Mg^2+^, and 5′-^32^P-mitochondrial-tRNA^Ile^. Time courses correspond to reactions measured at (1 – 180 min of total reaction) and reaction products were resolved by TLC. **(C)** Single turnover reaction curves for AcaTLP2 (top) and MxTLP (bottom) with 0.1 mM ATP, 5′-^32^P-mitochondrial-tRNA^Ile^ in the presence of 0.25 (red ●), 0.50 (blue ■), 1 (green ♦), 2 mM (open Δ) Mg^2+^. Total reaction products (App*C and Ap*C) were quantified and observed rates (k_obs_) were determined by fits to eq. 1, with each time course measured in at least two independent assays.

**Table 2:** Rates of adenylylation in the presence of varied [Mg^2+^]*^a^*.

| **$[Mg^{2+}]$**<br>**(mM)** | **AcaTLP2**<br>**$k_{obs}$ (min<sup>-1</sup>)** | **MxTLP**<br>**$k_{obs}$ (min<sup>-1</sup>)** |
| --- | --- | --- |
| 0.25 | 0.13 ± 0.01 | 0.017 ± 0.001 |
| 0.50 | 0.22 ± 0.01 | 0.018 ± 0.004 |
| 1.0 | 0.43 ± 0.05 | 0.020 ± 0.001 |
| 2.0 | 0.70 ± 0.05 | 0.032 ± 0.002 |

<sup>a</sup> Rates are averages of at least two independent measurements at each concentration

MxTLP and AcaTLP2 both exhibited modestly increased rates of 5’-adenylylation as the concentration of Mg^2+^ increased (**Table 2**). However, there was a distinct difference in the overall reaction amplitude between the two enzymes, with AcaTLP2 exhibiting consistently high endpoints (∼60%) at all tested concentrations of Mg^2+^, while MxTLP reaction endpoints increased across the concentration range (**Figure 4C**). Since the single turnover endpoint is an indicator of the maximal fraction of substrate that is convertible to product, but the same substrate was used in all of these assays, these data suggest that MxTLP acts more effectively on a specific RNA population, while AcaTLP2 is able to convert a higher fraction of the total RNA substrate pool. Given the known role of Mg^2+^ in tRNA stability(*24*), it is possible that MxTLP requires generally greater structural stability in its RNA substrates than AcaTLP2, which is more prevalent at higher [Mg^2+^] in the reactions. This distinct behavior of AcaTLP2 is again consistent with the recently observed catalytic flexibility of this enzyme (*22*). Whether this apparent aspect of substrate recognition is related to some characteristic of either enzyme’s physiological substrates remains unknown, since these substrates have not been identified.

We similarly measured single turnover rates of adenylylation catalyzed by AcaTLP2 and MxTLP in the presence of 2 mM Mn^2+^, Co^2+^, Ni^2+^, and Ca^2+^ (**Figure S1**). With AcaTLP2, the fastest rate of adenylylation occurs in Mn^2+^, with Co^2+^ and Mg^2+^ able to support this reaction at ∼50% slower rates (**Table 3**, **Figure 5**). With MxTLP, however, while Mg^2+^ and Mn^2+^ support adenylylation at similarly high rates, the observed rates in Co^2+^ and Ni^2+^ are ∼100 and ∼3000-fold slower, respectively. The decreased fitness of AcaTLP2 in Mg^2+^ compared to Mn^2+^ may be related to its apparently more relaxed base-pair selection fidelity, as evident from the significant amount of non-templated A_+1_-addition activity observed in these assays and discussed more below. The very slow product formation observed for 5’-adenylyation in Co^2+^ (for MxTLP), Ni^2+^ and Ca^2+^ required the use of the method of linear initial rates (eq. 2) to estimate k_obs_ for enzyme activity. Interestingly, for AcaTLP2, the difference in rate between the fastest (Mn^2+^) 5’-activation reaction and the Ni^2+^ reaction is only about 100-fold, whereas MxTLP activity in Ni^2+^ is ∼3,000-fold weaker than the fastest rate observed in Mg^2+^. Again, this suggests a relatively more flexible active site chemistry for AcaTLP2 than MxTLP. For Ca^2+^, the k_obs_ measured under the adenylyation assay conditions were extremely low for both enzymes, which was surprising since 5’-adenylylated intermediate had been readily detectable in the previous assays containing 2 mM Ca^2+^ (see **Figures 2C and 2D**). A similar discrepancy was observed between the very slow adenylylation rate of MxTLP in Co^2+^, despite high levels of product formation observed with this metal in Figure 2D. This phenomenon is explored further below.

**FIGURE 5.**
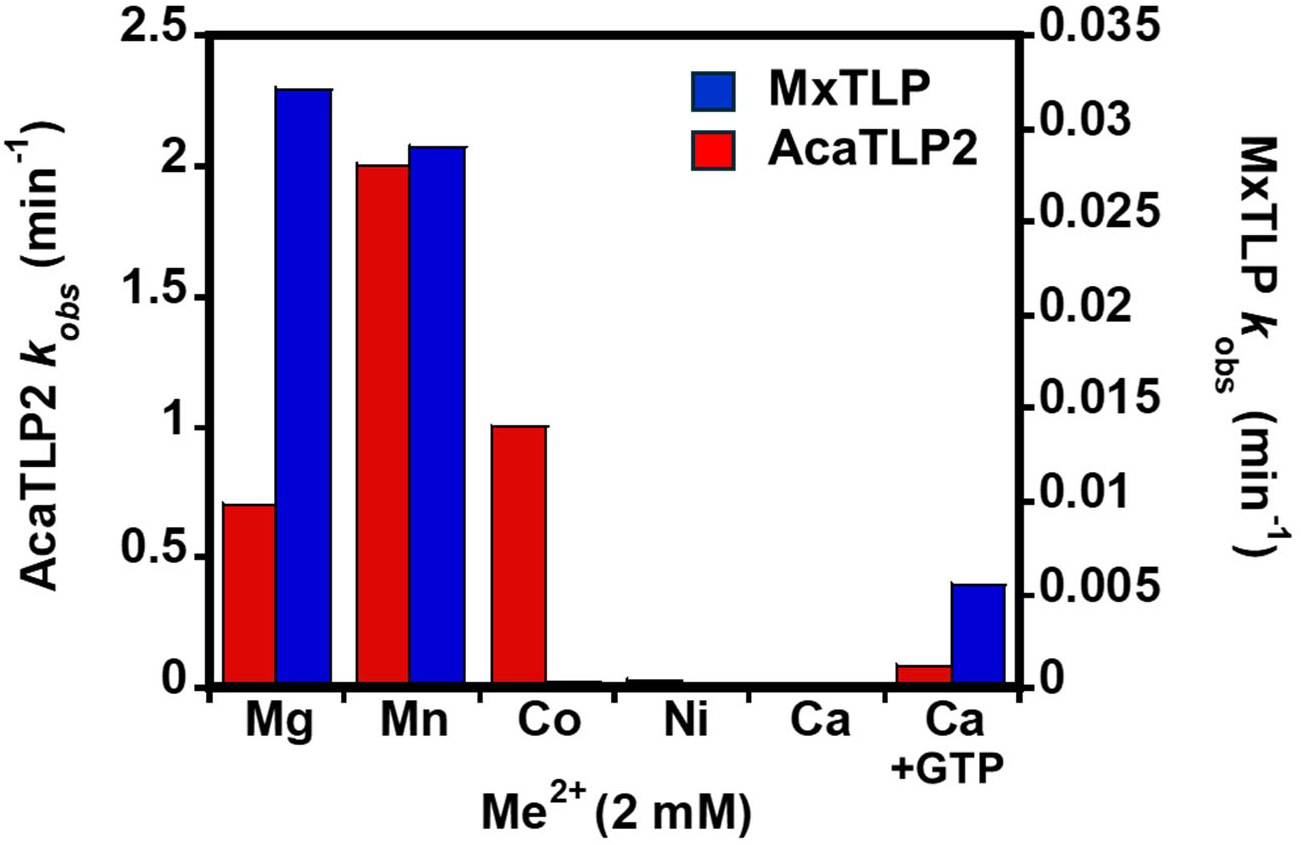
AcaTLP2 and MxTLP exhibit different patterns of cation-dependent 5’-adenylylation activity. Observed rates for 5’-adenylylation measured at 2 mM of each indicated metal using the assays described in Figure 4 are plotted for AcaTLP2 (red) and MxTLP (blue); note differences in scale on Y-axis for each enzyme. All reactions contained 0.1 mM ATP, except for reactions labeled Ca+GTP, which also contain 1 mM GTP.

**Table 3:** Rates of adenylylation in different divalent metals.

| **Metal**<br>**(2 mM)** | **AcaTLP2**<br>***k<sub>obs</sub>* (min<sup>-1</sup>)** | **MxTLP**<br>***k<sub>obs</sub>* (min<sup>-1</sup>)** |
| --- | --- | --- |
| Mg<sup>2+</sup> | 0.70 ± 0.05 | 0.032 ± 0.002 |
| Mn<sup>2+</sup> | 2.0 ± 0.2 | 0.029 ± 0.001 |
| Co<sup>2+</sup> | 1.0 ± 0.2 | 0.00021 ± 0.00001<sup>a</sup> |
| Ni<sup>2+</sup> | 0.027 ± 0.004<sup>a</sup> | 0.00001 ± 0.000002<sup>a</sup> |
| Ca<sup>2+</sup> | 0.000054 ± 0.000004<sup>a</sup> | 0.00006 ± 0.000005<sup>a</sup> |
| Ca<sup>2+</sup><br>(+GTP) | 0.083 ± 0.008 | 0.0055 ± 0.0001<sup>a</sup> |

<sup>a</sup> *k<sub>obs</sub>* was estimated using the method of linear initial rates (eq.2) due to low observed formation of product in assays

### Evidence for a pre-assembled active site containing both NTPs for activation and nucleotidyl transfer during 5’-adenylylation

To identify the source of the discrepancy between the amount of 5’-adenylate observed in the initial assays of each enzyme with Ca^2+^ (**Figure 2**) compared to the subsequent kinetic assays where very little 5’-adenylylation was observed in the same concentration of Ca^2+^ (**Figure S1**), we considered that the only difference between the two assays was the presence of both ATP and GTP nucleotides in Figure 2, vs. only ATP in Figure S1. It was not immediately clear why the presence of the GTP that would be used for the second step of the TLP reaction (nucleotidyl transfer) would impact the ability of the enzyme to carry out adenylylation since there is no obvious mechanistic role for the added nucleotide in the activation step of 3’-5’ addition (**Figure 1A**). To assess the role of other NTPs on the efficiency of the 5’-adenylylation step, we tested the effect of each of the other NTPs on the observed rates of this reaction in Ca^2+^ with AcaTLP2 and MxTLP (**Figure 6**).

**FIGURE 6.**
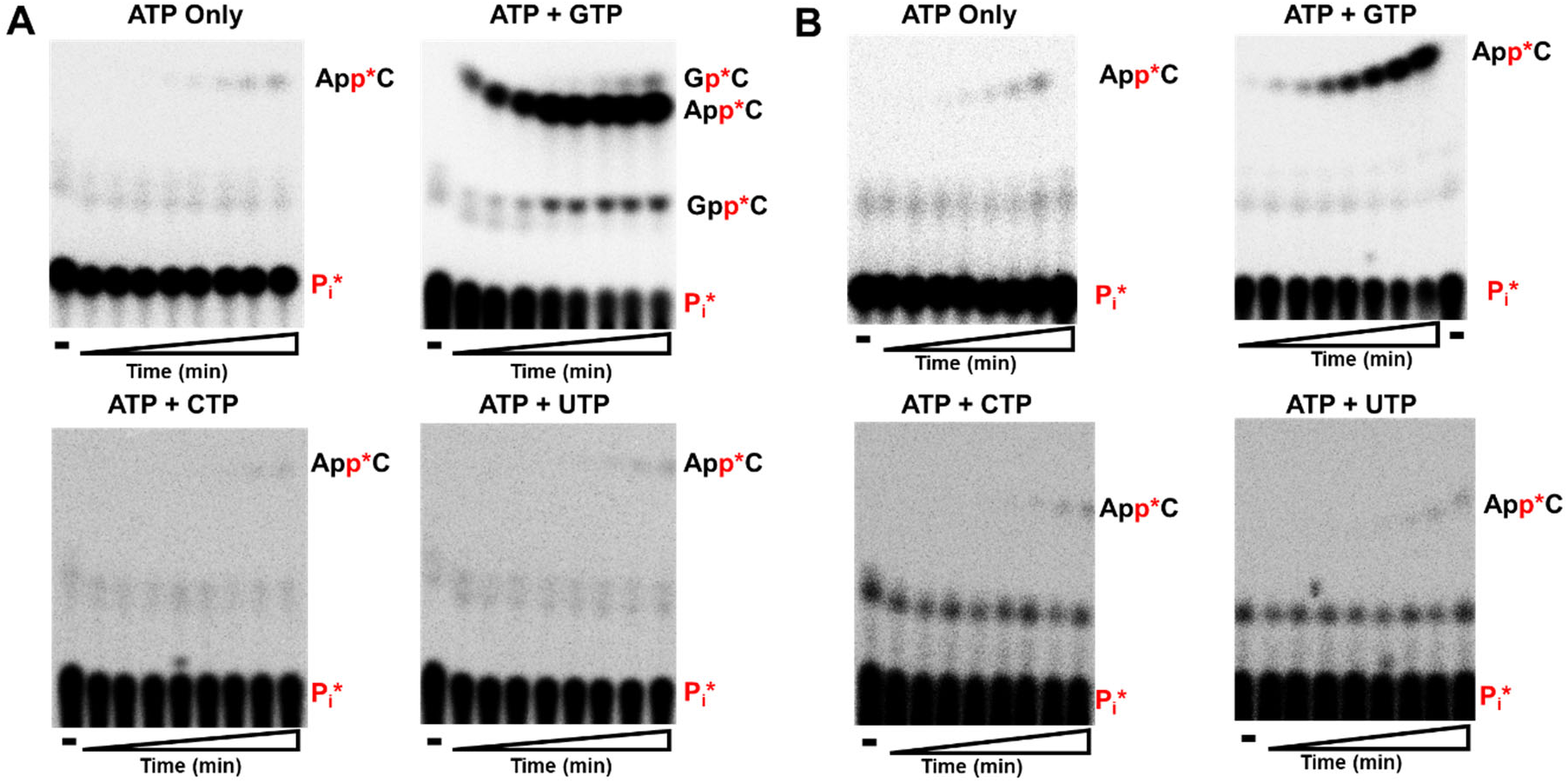
Ca^2+^ primarily allows only the adenylation step to occur with the correct Watson-Crick nucleotide. Single-turnover adenylation reactions containing **(A)** AcaTLP2 (10 µM) or **(B)** MxTLP (10 µM), and 0.1 mM ATP, 2 mM Ca^2+^, 5′-^32^P-mitochondrial-tRNA^Ile^. To each reaction, either no additional NTPs (top left), 1 mM GTP (top right), 1 mM CTP (bottom left), or 1 mM UTP (bottom right) is added prior to enzyme addition. Reaction products were resolved using silica TLC and identity of reaction products are shown on the right side of each panel.

In the presence of 2 mM Ca^2+^ and 1 mM GTP (the correct Watson-Crick NTP to base pair with C_72_ in this substrate tRNA, see Figure 2A), the rate of 5’-adenylylation increased dramatically from the measured rate in ATP only for both enzymes (**Table 3**). At the longest reaction times with AcaTLP2, we also observed a small amount of what is likely the 5’-guanylylated product as explained previously, and even a very small amount of the G_+1_ addition product (**Figure 6A**). This increased rate of 5’-activation was selective for the ability of the added NTP to form a Watson-Crick base pair with the C_72_ templating nucleotide on the RNA, since inclusion of either 1 mM CTP or UTP (lower panels) appear similar to the ATP-only reactions. With MxTLP, similar overall results were observed; despite the overall slower k_obs_ values measured for this enzyme, measurable rates of adenylylation were only obtained in reactions containing both ATP and GTP (but not any other NTP combinations) (**Figure 6B** and **Table 3**).

To test whether the stimulation of step 1 by the presence of the NTP to be used in the subsequent nucleotidyl transfer step can be observed with other base-pairing combinations, we created a different RNA substrate (5′-[^32^P]-tRNA^Leu^ U_+3_), for which the templating nucleotide is a G_71_ residue; and therefore CTP is the correct Watson-Crick base paired NTP for nucleotidyl transfer (**Figure S2**). We note that this substrate also theoretically allows for UTP to be incorporated as a U_+2_:G_71_ wobble base pair, allowing us to test whether the stimulatory effect is dependent on recognition of some aspect of Watson-Crick base pairing or is more generally due to a stabilizing effect of hydrogen bonding with the incoming NTP in the enzyme active site. The overall rates of adenylation with this substrate are generally lower than with the mt-tRNA^Ile^ tested earlier, such that measurable activity was only observed with AcaTLP2. Nonetheless, these assays showed that the presence of the correct Watson-Crick base pairing CTP modestly stimulates the efficiency of adenylylation relative to the ATP-only reaction. Interestingly, the presence of UTP inhibited activity with this substrate completely, while the presence of GTP did not further enhance the efficiency of 5’-activation.

## Discussion

In this study, we used biochemical approaches to investigate the metal ion dependence for several members of the 3′-5′ polymerase family. Prior to this study, the activity of all TLPs had been characterized in a standard buffer containing 10 mM Mg^2+^, based on the presumption that this is the physiologically relevant metal ion for 3’-5’ polymerase activity. We demonstrated that, much like canonical two metal ion-dependent polymerases, 3’-5’ polymerases can utilize other divalent cations for catalysis (**Figure 2**). Using two TLP enzymes from different domains of life, we demonstrated that Mn^2+^, Mg^2+^, Co^2+^ and Ni^2+^ are all able to support both 5’-activation and nucleotidyl transfer steps of 3’-5’ polymerase activity, albeit in distinct ways and with different overall efficiencies that reflect enzyme-specific preferences (**Figures 2-5**). Ca^2+^, on the other hand, is only able to support the chemistry of adenylylation (step 1) and does not allow the enzyme to catalyze nucleotidyl transfer (step 2) to any appreciable extent (**Figure 2**). Using Ca^2+^, we identified an unexpected role of the incoming base-paired NTP for nucleotidyl transfer on the previous (adenylylation) step of the reaction, suggesting this nucleotide may form a pre-existing base pair in the active site, even before it is actually needed for catalysis (**Figure 6**). These data indicate that rearrangement of nucleotides around the two metal ions during the transition from 5’-end activation to nucleotidyl transfer is an important mechanistic step that remains to be fully understood but must be able to accommodate the presence of multiple different NTPs for efficient catalysis.

Crystal structures of members of the 3′-5′ polymerase family have provided important information about the mechanism of Thg1/TLP catalysis (*12, 13, 18, 20*). In all 3’-5’ polymerase structures to date, at least two metals have been observed in the active site, with some structures showing the presence of a third metal, which appears to play a role in positioning the incoming NTP for nucleotidyl transfer (2) with 5’-triphosphorylated RNA (*20*). The similar arrangement of Metal A and Metal B relative to the attacking nucleophile and scissile phosphate in Thg1/TLP structures supported the assignment of analogous roles to the known roles of these metals in canonical 5’-3’ polymerases (**Figure 1**) (*25*). However, members of the Thg1/TLP family are more complex, given that they are capable of catalyzing two distinct reactions (5’-adenylylation and nucleotidyl transfer) with the same two metal ion active site. In this way, 3’-5’ polymerases also exhibit some similarity to ligases, which also create 5’-adenylylates from 5’-monophosphorylated RNA ends prior to joining (“sealing”) their substrates (*26, 27*). The characterization of TLP metal ion dependence here revealed features that are similar to some aspects of metal ion usage by both polymerases and ligases, suggesting that 3’-5’ polymerases represent somewhat of a “hybrid” mechanism. For nucleotidyl transfer, AcaTLP2 and MxTLP are strongly active in Mg^2+^ and Mn^2+^, with variable amounts of activity in Co^2+^ and Ni^2+^, and no activity in Ca^2+^ (**Figure 2**). This is very similar to the pattern of metal dependence commonly observed for many DNA and RNA polymerases and correlates with the expected properties of each metal to adopt the appropriate coordination geometry and exhibit the optimal pKa to effectively activate the 3’-OH nucleophile for attack on the scissile phosphate (*28, 29*). Yet, for adenylylation, Ca^2+^ also supports readily detectable activity of both tested TLPs, which is also commonly observed in ligases that can use Ca^2+^ for this first 5’-adenylylation step during their end sealing reaction. Moreover, the different patterns of activity do not always follow patterns that can be determined from their functional properties. While Ni^2+^ and Co^2+^ show similar abilities to weakly support adenylation by MxTLP that are consistent with their similar ionic radii (0.83 Å vs. 0.89 Å, respectively) and pKa values (10.0 vs. 10.6, respectively), there is a much larger difference in adenylation rate observed between these same two metals with AcaTLP2 (**Figure 5**). Thus, understanding the unique molecular environment of each enzyme with different metals will be needed to understand these distinct preferences.

Like with 5’-3’ polymerases, it is likely that Mg^2+^ is the physiological activator of Thg1/TLPs due to its overall abundance, its ability to universally activate these polymerases, and its ability to support high fidelity polymerase activity. Although Mn^2+^ and Co^2+^ also support activity for some canonical 5’-3’ DNA polymerases, both metals have been classified as mutagens because of their reduced fidelity of nucleotide incorporation. Although we did not explicitly test metal-dependent fidelity of NTP addition in this study, evidence for similar mutagenic behavior of Mn^2+^ and Co^2+^ with TLPs was observed when looking at the non-Watson-Crick addition of A_+1_ to mt-tRNA^Ile^ (**Figure 4** and **Figure S1**). Although AcaTLP2 readily forms the A_+1_pC product even in Mg^2+^ (**Figure 4B**), this non-Watson-Crick product corresponds to nearly all of the observed product in Mn^2+^ and is formed at much earlier time points in the presence of Co^2+^ compared to in Mg^2+^ (**Figure S1**). With MxTLP, there is a similar effect, with no detectable non-Watson-Crick addition of A_+1_ in Mg^2+^, but this addition is readily seen in Mn^2+^ (**Figure S1**). It is intriguing to speculate that this ready ability of AcaTLP2 to “misincorporate” is related to its biological function, which is currently not known, and the data here could suggest that MxTLP is more stringent than AcaTLP2 under normal cellular conditions.

While we were able to quantify the impact of metal identity and concentration on the isolated 5’-adenylation step of 3’-5’ polymerases here, our data clearly demonstrate that understanding the impact of metals on the nucleotidyl transfer reaction would also be insightful. Unfortunately, the [γ-^32^P]-nucleotide we used previously to generate appropriately labeled RNA substrates that allowed us to isolate and measure rates for nucleotidyl transfer during 3’-5’ addition is no longer commercially available (*8, 14, 23, 30*). Thus, directly assessing the impact of metals on these rates will require development of alternative kinetic assays. Nonetheless, these data highlight many yet unanswered questions about the transitions between different steps of the 3’-5’ polymerase reaction. In this case, it is important to consider rearrangements of each phosphate-containing substrate around each metal (i.e., note the interaction of the 5’-α-phosphate of ATP with Me_A_ during adenylylation followed by an inferred interaction of this same phosphate with Me_B_ during nucleotidyl transfer (1) as shown in **Figure 1B**, and other similar apparent movements of functional groups that must occur during catalysis). Our demonstration that the incoming base-paired NTP occupies the active site even during the earlier adenylylation step (Figure 6) suggests that all three metals may be present throughout catalysis and adds an additional layer of complexity to understanding these transitions. In Thg1/TLP structures in which the third metal ion (Me_C_) has been visualized, this metal has been implicated kinetically and structurally in the nucleotidyl transfer step (*12, 14, 20*). Given the ability of Ca^2+^ to facilitate adenylylation but not nucleotidyl transfer, it is also possible that the ionic radius of Ca^2+^ (1.11Å) does not allow rearrangement of the active site or function appropriately in this third metal ion binding site to permit nucleotidyl transfer to occur.

The possibility that the Thg1/TLP active site is sensitive to the presence of a correct base-paired NTP for nucleotidyl transfer during the 5’-adenylylation step of the reaction had been suspected previously. In studies with human Thg1, two conserved amino acids (His152 and Lys187) were implicated in base pair recognition, yet their major kinetic impact was observed on the adenylylation step (*23*). Based on the results of alanine substitutions, His152 was suggested to participate in a stabilizing interaction with the incoming non-Watson-Crick GTP. However, His152 and Lys187 are conserved eukaryotic amino acids in eukaryotic Thg1 and do not have direct counterparts in TLPs. The increased k_obs_ seen here for adenylylation in the presence of the next cognate Watson-Crick forming NTP provides the first direct evidence that both the ATP and the NTP to be incorporated are likely both present prior to and during the adenylylation step.

Furthermore, when an incorrect NTP is present, enzyme activity is reduced, suggesting that the inability to create a Watson-Crick base pair could impact the rate of adenylylation through an altered conformation of the active site. Interestingly, the presence of the NTP capable of forming a wobble (G●U) base pair did not increase the rate, despite the thermodynamic stability and ubiquity of these base pairs in RNA structures, supporting the idea that the incoming nucleotide has to form a true Watson-Crick base pair. This observation highlights the need for additional structures of Thg1/TLP enzyme complexes to elucidate all of the NTP binding modes and rearrangements around the metal ions that likely occur during catalysis.

## ASSOCIATED CONTENT

### Supporting Information

The following files are available free of charge.

Figure S1: Time courses of adenylylation activity in Mn^2+,^ Co^2+^, Ni^2+^ and Ca^2+^ (PDF) Figure S2: Time courses of adenylylation activity with RNA substrate templating for CTP addition (PDF)

## AUTHOR INFORMATION

### Author Contributions

The manuscript was written through contributions of all authors. All authors have given approval to the final version of the manuscript.

## ACKNOWLEDGMENT

The authors would like to thank the members of the Jackman lab for their advice during the experimental procedures and insightful conversations for manuscript preparation.

## ABBREVIATIONS

Thg1: tRNA^His^ guanylyltransferase
TLPs: Thg1-like proteins
PNK: polynucleotide kinase
RNase A: Ribonuclease A
CI(A)P: Calf intestinal (alkaline) phosphatase
MxTLP: *M. xanthus* TLP
AcaTLP2: *A. castellanii* TLP
MsTLP: *M. smithii* TLP
BtTLP: *B. thuringiensis* TLP
DdiTLP4: *D. discoideum* TLP4
HEPES: 4-2-hydroxyethyl-1-piperazineethanesulfonic acid
EDTA: ethylenediaminetetraacetic acid
DTT: dithiothreitol
TLC: thin layer chromatography;
mt-tRNA: mitochondrial tRNA.

