## Supplemental Figures for "Metal ion requirement for catalysis by 3′-5′ RNA polymerases"

#### SUPPLEMENTAL FIGURE LEGENDS

**Supplemental Figure 1.** Single-turnover adenylation reactions containing **(A)** 10 μM AcaTLP2 or **(B)** 10 μM MxTLP and 0.1 mM ATP, 5′-$^{32}$P-mitochondrial-tRNA$^{Ile}$, and either 2 mM $Mn^{2+}$ (top left), 2 mM $Co^{2+}$ (top right), 2 mM $Ni^{2+}$ (bottom left), or 2 mM $Ca^{2+}$ (bottom right). Reaction products were resolved using silica TLC and identity of reaction products are shown on the right side of each panel. The $k_{obs}$ value determined from eq. 1 are shown in Table 3.

**Supplemental Figure 2.** **(A)** Schematic of the 3′-5′ adenylylation reaction performed by a standard TLP to activate the 5′-end of 5′-$^{32}$P-mitochondrial-tRNA$^{Leu}$ from *D. discoideum*. Below each tRNA is the radiolabeled product formed when the transcript is treated with RNase A and CIP, which can be visualized via TLC. **(B)** Single-turnover adenylation reactions containing 10 μM of AcaTLP2, 0.1 mM ATP, 2 mM $Ca^{2+}$, 5′-$^{32}$P-mitochondrial-tRNA$^{Leu}$, and either 1 mM GTP (top right), 1 mM CTP (bottom left), or 1 mM UTP (bottom right). Reaction products were resolved using silica TLC and identity of reaction products are shown on the right side of each panel.

#### Supplementary Figure 1

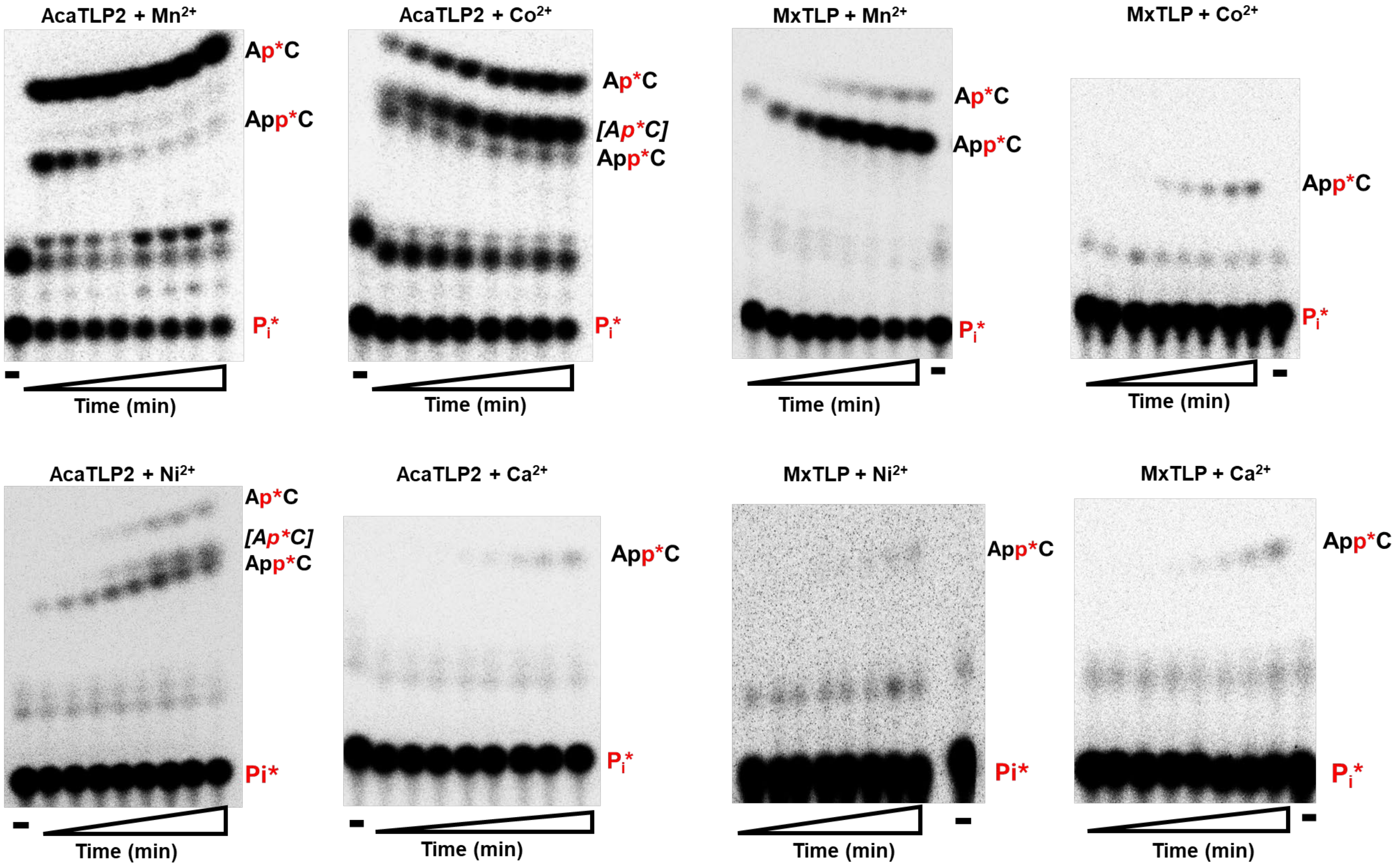

### Supplementary Figure 2

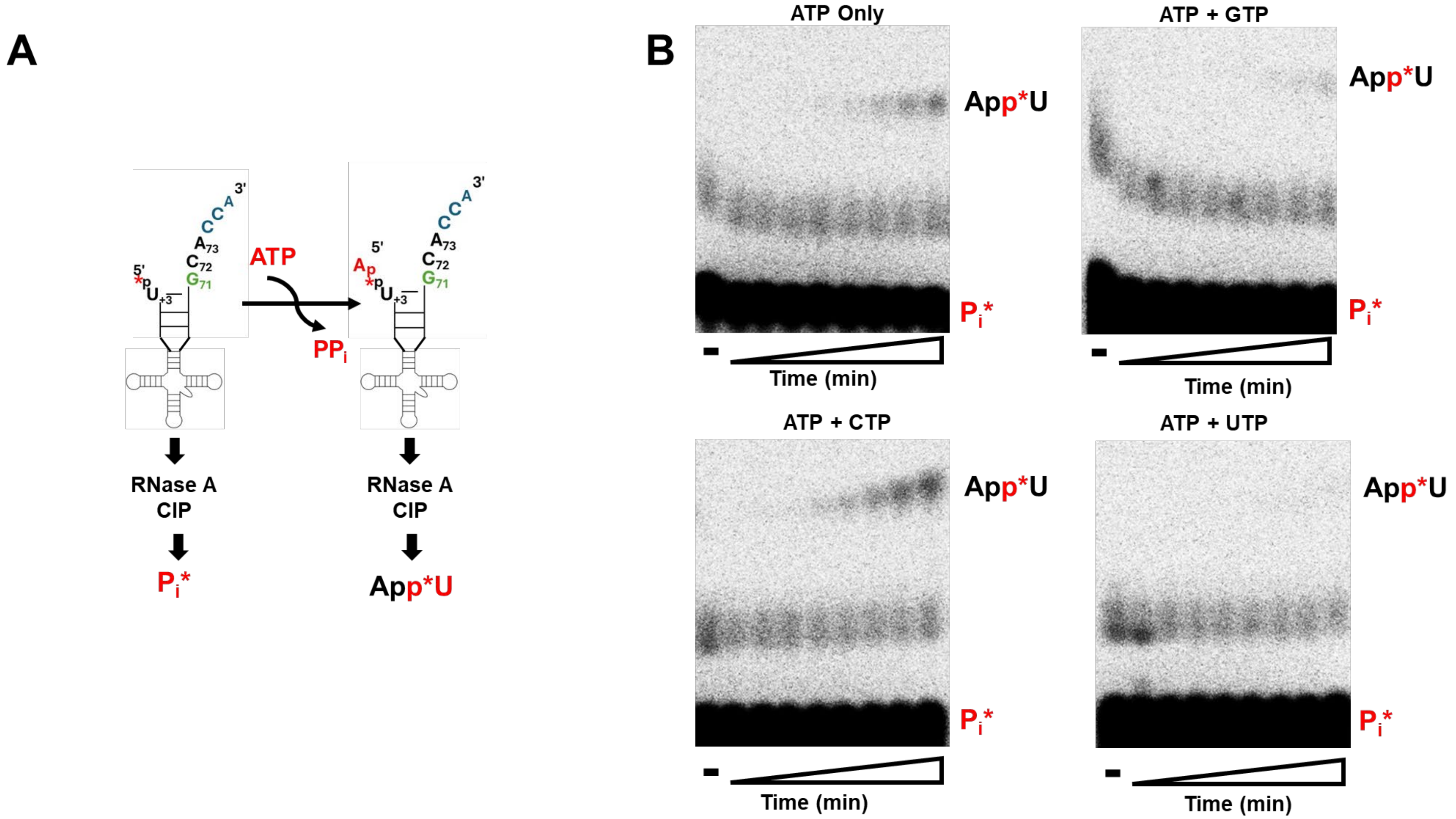
